# Metformin in nucleus accumbens core reduces cue-induced cocaine seeking in male and female rats

**DOI:** 10.1101/2021.09.05.458975

**Authors:** Amy Chan, Alexis Willard, Sarah Mulloy, Noor Ibrahim, Allegra Sciaccotta, Mark Schonfeld, Sade M Spencer

## Abstract

This study investigated the potential therapeutic effects of the FDA-approved drug metformin on cue-induced reinstatement of cocaine seeking. Metformin (dimethyl-biguanide) is a first-line treatment for type II diabetes that, among other mechanisms, is involved in the activation of adenosine monophosphate activated protein kinase (AMPK). Cocaine self-administration and extinction is associated with decreased levels of phosphorylated AMPK within the nucleus accumbens core (NAcore). Previously it was shown that increasing AMPK activity in the NAcore decreased cue-induced reinstatement of cocaine seeking. Decreasing AMPK activity produced the opposite effect. The goal of the present study was to determine if metformin in the NAcore reduces cue-induced cocaine seeking in adult male and female Sprague Dawley rats. Rats were trained to self-administer cocaine followed by extinction prior to cue-induced reinstatement trials. Metformin microinjected in the NAcore attenuated cue-induced reinstatement in male and female rats. Importantly, metformin’s effects on cocaine seeking were not due to a general depression of spontaneous locomotor activity. In female rats, metformin’s effects did generalize to a reduction in cue-induced reinstatement of sucrose seeking. These data support a potential role for metformin as a pharmacotherapy for cocaine use disorder, but warrant caution given the potential for metformin’s effects to generalize to a natural reward in female rats.

## Introduction

Despite decades of research, cocaine use disorder remains a significant public health concern in the United States. In the 2018 National Survey on Drug Use and Health, 5.5 million people aged 12 and older (approximately 2% of the population) reported past month cocaine use.^1^ Further highlighting the urgency of the problem, frequencies of cocaine-involved overdose deaths are on the rise.^2^ Unfortunately, there are still no FDA-approved pharmacotherapies for the treatment of cocaine use disorder. Even when abstinence is obtained, relapse rates for this chronic disease remain high.^3^ Males and females show behavioral differences in substance use dependent on biological influences including sex hormones.^4^ In addition, there may be sex differences in response to pharmacological treatments for substance use disorders.^5^ Accordingly, there is a well-defined need to identify and characterize novel pharmacotherapies for preventing cocaine relapse using male and female preclinical subjects.

Adenosine monophosphate activated protein kinase (AMPK) is a heterotrimeric serine/threonine protein kinase that acts as a cellular energy sensor monitoring the ratio of AMP to ATP in the cell.^6^ AMPK agonists are used to treat conditions including type II diabetes, cancer, and cardiovascular disease.^7^ Nascent literature suggests that AMPK can regulate drug-associated behaviors including locomotor sensitization to alcohol and cocaine.^8^ One recent study demonstrated that microinjection of the AMP analog 5-aminoimidazole-4-carboxamide ribonucleoside (AICAR) in nucleus accumbens core (NAcore) reduces cue-induced reinstatement in a rat model of cocaine seeking.^9^ However, the translational utility of AICAR is limited because it has poor pharmacokinetic properties and lacks oral bioavailability.^7,10^ Moreover, high doses of AICAR are associated with numerous deleterious side effects including bradycardia and other metabolic problems.^7^ In contrast, metformin (dimethyl-biguanide), the most commonly prescribed FDA-approved treatment for type II diabetes, is a safe and widely-used drug that can activate AMPK among other mechanisms of action.^11^ In preclinical studies, metformin was shown to alleviate nicotine withdrawal-induced anxiety-related behaviors, attenuate the development of morphine analgesic tolerance, and reduce opioid withdrawal symptoms.^12,13^ No studies have examined the efficacy of metformin, an arguably more translationally relevant AMPK activator compared to AICAR, to reduce the reinstatement of cocaine seeking behavior. Importantly, metformin readily crosses the blood-brain barrier and has an absolute oral bioavailability of 40-60% in humans.^14^

The present study assessed the ability of intracranial microinjections of metformin in the NAcore to attenuate the reinstatement of drug-seeking behavior elicited by response contingent cocaine conditioned cues in male and female rats. Both the nucleus accumbens (NAc) core and shell are implicated in drug seeking responses,^15^ but NAcore and NAc shell display dissociable roles in responding to cocaine-paired conditioned reinforcers.^16^ Phosphorylated AMPK is down regulated in the NAcore following extinction from cocaine self-administration while levels remain unchanged in the NAc shell.^9^ Such regionally selective modulation of the AMPK system by cocaine provides additional rationale for our present focus on the NAcore.^9,17^ We resolved that the effects of intracranial metformin on cocaine seeking were independent of effects on locomotor activity. We also determined whether the effects of metformin in NAcore were selective for cue-induced cocaine seeking or generalized to cue-induced seeking for sucrose, a natural reward. These data establish a foundation for further pursuing metformin as a novel pharmacotherapy for cocaine relapse.

## Materials and Methods

### Subjects

137 male and female Sprague Dawley rats (225-275g on arrival) were purchased from Envigo, and 109 subjects were included in the final dataset. Rats were pair-housed in individually ventilated cages in a temperature (70-71°F) and humidity (38-46%) controlled holding facility. Standard chow and water were available *ad libitum* unless otherwise specified. Animals were kept on a 14 /10 light-dark cycle with all experiments conducted during the light phase. Procedures were preapproved by the Institutional Animal Care and Use Committee (IACUC) of the University of Minnesota and follow National Institute of Health (NIH) guidelines.

### Surgery

Rats (250-300g at time of surgery) were anesthetized with vaporized isoflurane (2-5%). Rats were implanted with intravenous jugular catheters and bilateral guide cannula (Plastics One) to NAcore: ±1.8 mm ML, +1.5 mm AP, −5.5 mm DV and ±1.6 mm ML, +2.3 mm AP, −5.5 mm DV at a 0° angle. Postoperative analgesia was provided for 72 hours post-surgery (carprofen, 5 mg/kg i.p.). Catheter patency was confirmed prior to initiating cocaine self-administration with intravenous administration of 0.05-0.1 mL xylazine (5 mg/mL).

### Drugs

Cocaine hydrochloride was purchased from Boynton Pharmacy at University of Minnesota. Cocaine solution (4 mg/ml) was prepared in sterile 0.9% saline and delivered at a volume of 0.05 ml/infusion. On average male and female rats self-administered cocaine doses of 0.59 and 0.74 mg/kg/infusion, respectively over the course of self-administration based on body weight disparity (Figure S1). Metformin hydrochloride (ApexBio, Sigma-Aldrich), 1,1-dimethylbiguanide) was prepared at 250 μg/μl in sterile water, and 125 μg/hemisphere was microinjected into NAcore. The metformin dose was based on published studies of intracerebroventricular (ICV) administration in rats using doses ranging from 3 μg to 1 mg.^18–20^ Saline or metformin was microinjected in NAcore with brain dissections collected after 1 hour in a subset of animals used for Western blotting. As expected, metformin increased levels of phospho-AMPK and showed a trend toward decreased phospho-ERK consistent with AMPK activation (Figure S2).

### Cocaine self-administration, extinction, and cue-induced reinstatement

Training was completed in standard operant boxes (MedAssociates). Rats underwent one 2-hour session of operant training for food pellets in the absence of cues prior to beginning daily 2-hour cocaine self-administration sessions. Overnight food restriction preceded the food training session. Rats self-administered cocaine on a fixed ratio 1 (FR1) reinforcement schedule. Each cocaine infusion was paired with a discrete white light and tone dual stimulus. Inactive lever presses were recorded but had no consequences. Rats were required to reach a criterion of 10 days of self-administration with ≥10 cocaine infusions. During extinction (minimum 7 days) pressing the formerly active lever no longer resulted in delivery of cocaine or cues. Extinction required reducing active lever pressing to ≤30% of the self-administration mean. During cue-induced reinstatement active lever pressing resulted in presentation of drug-paired cues but no drug delivery during a 2-hour session.

### Sucrose self-administration, extinction, and cue-induced reinstatement

The protocol for sucrose self-administration was analogous to that used for cocaine, described above. Rather than cocaine infusions, sucrose pellets (45mg, BioServ) were paired with a discrete light and tone cue. Rats were trained to self-administer sucrose pellets for at least 5 days on an FR1 reinforcement schedule and then progressed to 5 days on an FR3 reinforcement schedule with acquisition criterion of ≥10 sucrose pellets earned per session. Extinction and cue-induced reinstatement conditions were identical to cocaine self-administration.

### Locomotor Activity

Spontaneous locomotor activity was recorded with a USB webcam placed over the test chamber (60cm x 60cm square acrylic box) for 45 minutes. Videos were analyzed for total distance traveled using ezTrack video analysis software. Animals were given microinjections of saline or metformin to the NAcore immediately before or 1 hour prior to placement in test chamber.

### Microinjections and Histology

Metformin or 0.9% sterile saline was microinjected into NAcore 2 mm below guide cannula base. Microinjections (0.5 μl/side) occurred over 2 minutes, followed by 1 minute of diffusion time. The reinstatement session started immediately after microinjection. For within-subject comparisons of experimental treatment on cue-induced reinstatement (saline vs. metformin), rats underwent two reinstatement sessions separated by ≥2 extinction sessions. The order was randomized between subjects. An equivalent strategy was used to assess metformin’s effects on cocaine self-administration maintenance (maintenance day 7 and day 9-10) and extinction responding (extinction day 2 and day 4-5). For histology, animals were deeply anesthetized with sodium pentobarbital (80-100 mg/kg) and transcardially perfused with 1x phosphate buffered saline (PBS, in MilliQ H2O) followed by 1x Formalin in PBS. Coronal slices of NAcore (100 μm) were mounted on glass slides and stained with cresyl violet to validate cannula placement. Slides were imaged using an Olympus stereomicroscope (MVX10) and cannula placements were visually confirmed. Histological verification of microinjection placement for all experiments is shown in the supplementary materials (Figure S5, S6). Rats with off-target cannula were excluded from analysis. Successful bilateral and unilateral (due to cannula occlusion) microinjections were included in the final dataset after supplementary analysis revealed equivalent trends in the reinstatement effects in the subdivided data (Figure S6).

### Statistical Analysis

GraphPad Prism version 9.0.1 (San Diego, California USA) was used to perform statistical analysis. Data were analyzed using 1- or 2-way ANOVAs or using mixed-effects models as appropriate for each experiment (see supplementary materials for additional details). Tukey’s or Sidak’s *post-hoc* tests were applied for multiple comparisons and *p*<0.05 was considered statistically significant. Data are presented as the mean ± standard error of the mean (SEM). In the main text, data for each sex is presented separately, but direct comparisons between males and females can be found in the supplementary figures (Figures S1, S3, S4 and S5).

## Results

### Equivalent cocaine self-administration in male and female Sprague Dawley rats

The experimental timeline is depicted in Figure 1A. Male and female rats were trained to self-administer cocaine and then extinguished for 7-14 days (Figure 1B, C). Rats self-administered similar numbers of infusions per day over the course of the 10 sessions (Figure 1D), and lever pressing did not differ between sexes (Figure S3A, B). When comparing the discrimination index of males and females using mixed-effects analysis there was a main effect of session [F(5.948,99.30)=4.363, p=0.0006], a main effect of sex [F(1,20)=5.621, p=0.0279], and a significant session by sex interaction [F(23,384)=3.120, p<0.0001] (Figure 1E). While male rats showed decreased discrimination during the extinction phase, female rats maintained preferential responding on the previously reinforced lever. Estrous cycle was monitored by vaginal cytology throughout the experiment. There was no effect of estrous cycle stage on active lever pressing (Figure 1F) or cocaine infusions (Figure 1G) during the self-administration phase. The analysis of day 1 extinction active lever pressing combined proestrus and estrus (follicular phase) compared to metestrus and diestrus (luteal phase) given the limited number of observations. Female rats in proestrus/estrus showed greater levels of extinction day 1 active lever pressing compared to metestrus/diestrus [Figure 1H; T(10)=2.663, p=0.0236], but no sex difference was observed.

**Figure 1.**
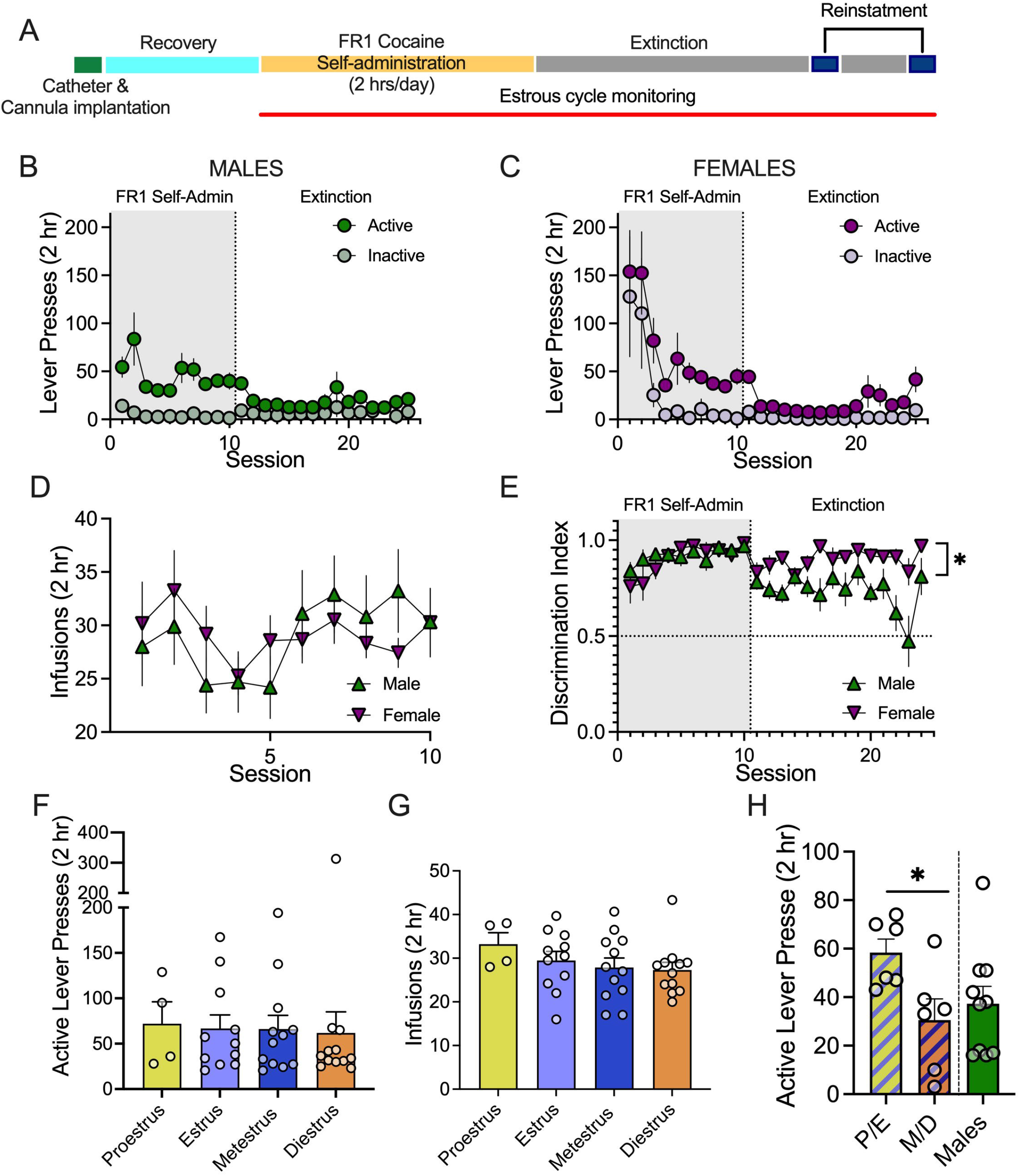
Cocaine self-administration in male and female Sprague Dawley rats. A. Experimental timeline for cocaine self-administration, extinction, and reinstatement. B. Active and inactive lever pressing in males. C. Active and inactive lever pressing in females. D. Daily cocaine infusions over the 10 days of self-administration at criteria did not differ between male and female rats. E. The discrimination index remained higher in female rats compared to males during the extinction phase, *p<0.05, n=10-12 per sex. F. Self-administration active lever pressing did not vary by estrous cycle stage. G. Cocaine infusions delivered during self-administration did not vary by estrous cycle stage. In F. and G. each data point represents the rat mean for each cycle stage. H. Active lever pressing on extinction day 1 was higher during the follicular phase (P/E = proestrus/estrus) compared to the luteal phase (M/D = metestrus/diestrus), *p<0.05, n=3-4 per phase. All data expressed as mean ± SEM.

### Metformin reduces cue-induced reinstatement of cocaine seeking

Cue-induced reinstatement of cocaine seeking was tested using a randomized within-subjects crossover design comparing saline and metformin microinjection in NAcore. There was a main effect of lever [F(1,18)=53.51, p<0.0001], a main effect of condition [F(1.494,26.89)=6.930, p=0.0013], and a significant interaction [F(2,36)=8.018, p=0.0013] for cue-induced reinstatement of cocaine seeking in males as assessed by two-way ANOVA (Figure 2A). Tukey’s multiple comparisons testing indicated a significant increase in active lever pressing during reinstatement compared to extinction after saline microinjection (p=0.0075). After metformin microinjection, reinstatement of active lever pressing was not significantly different from either extinction levels (p=0.1583) or saline pretreatment (p=0.2655) (Figure 2A). We interpreted this as a lack of reinstatement in the metformin condition. Cumulative lever responses were assessed during cue-induced reinstatement trials to determine if there were time course-dependent effects of metformin. In the male rats, analysis of active lever pressing showed a main effect of treatment [F(1,18)=5.021, p=0.0379], a main effect of time [F(1.498,26.96)=22.46, p<0.0001], and a significant treatment x time interaction [F(23,414)=5.006, p<0.0001] (Figure 2C). Metformin’s effects on drug seeking occurred after the first hour of reinstatement. When examined separately, active lever pressing during reinstatement did not differ significantly for saline versus metformin treatment in the first hour (p=.1518), but this effect emerged during the second hour (p=0.0207). Metformin treatment did not significantly modify inactive lever pressing during reinstatement (Figure 2E).

**Figure 2.**
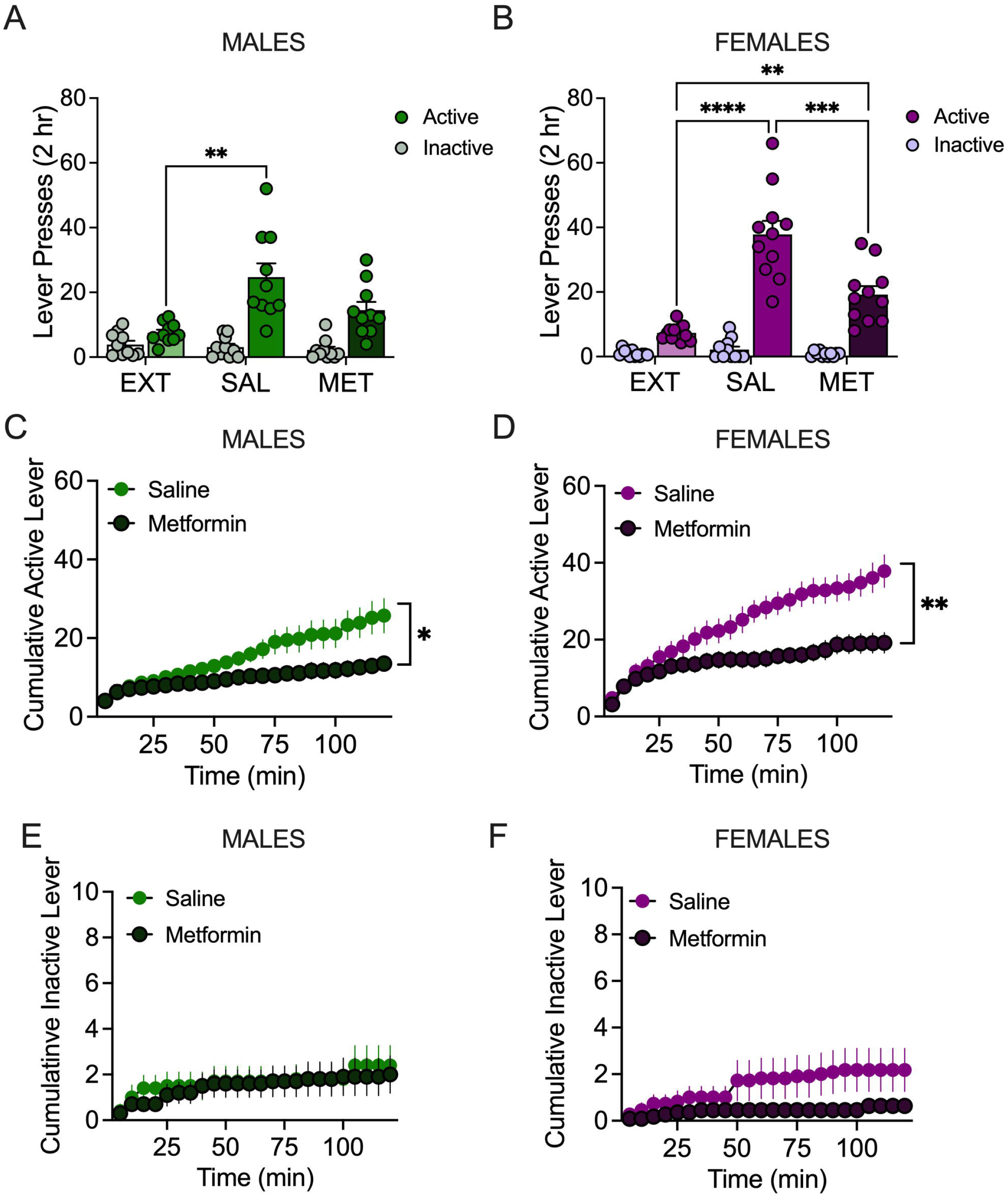
Effects of metformin on cue-induced reinstatement of cocaine seeking. A. Number of active and inactive lever presses during extinction (EXT) or induced by cocaine-associated cues during reinstatement in male rats, n= 10. **p<0.01 comparing saline (SAL) to extinction (MET = metformin). B. Number of active and inactive lever presses during extinction or induced by cocaine-associated cues during reinstatement in female rats, n= 12. ****p<0.0001 comparing saline to extinction; **p<0.01 comparing metformin to extinction or saline. C. Cumulative active lever pressing in 5-minute bins during cue-induced reinstatement in male rats. *p<0.05 comparing saline to metformin. D. Cumulative active lever pressing in 5-minute bins during cue-induced reinstatement in female rats.*p<0.01 comparing saline to metformin. E. Cumulative inactive lever pressing in 5minute bins during cue-induced reinstatement in male rats. F. Cumulative inactive lever pressing in 5-minute bins during cue-induced reinstatement in female rats. All data expressed as mean ± SEM.

**Figure 3.**
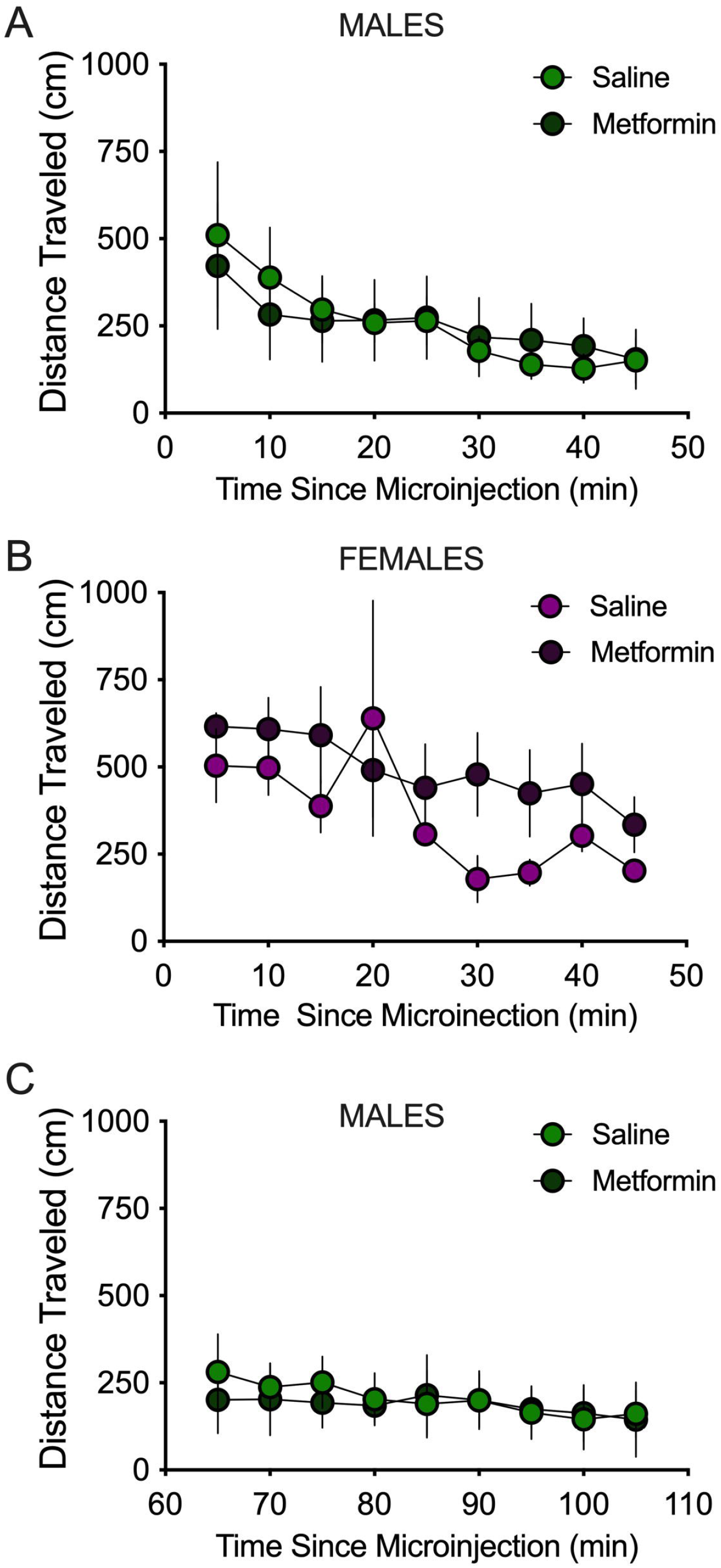
Locomotor activity. A. Locomotor activity in male rats immediately after microinjection of saline or metformin, n=5 per treatment. B. Locomotor activity in female rats immediately after microinjection of saline or metformin, n=5-7 per treatment. C. Locomotor activity in male rats following 1 hour pretreatment time after saline or metformin microinjection, n=7 per treatment. All data expressed as mean ± SEM.

In female rats, there was a main effect of lever [F(1,22)=99.36, p<0.0001], a main effect of condition [F(1.718,34.37)=42.39, p<0.0001], and a significant interaction [F(2,40)=36.07, p<0.0001] for cue-induced reinstatement of cocaine seeking (Figure 2B). Tukey’s multiple comparisons testing indicated a significant increase in active lever pressing during reinstatement compared to extinction after both saline (p<0.0001) and metformin microinjection (p=0.0050) indicating cue-induced reinstatement under both conditions (Figure 2B). Lever pressing was also significantly lower after metformin pretreatment compared to saline (p=0.0007) (Figure 2B). When time bin analysis was performed in the female rats, there was a main effect of treatment [F(1,20)=8.751, p=0.0078], a main effect of time [F(2.512,50.24)=73.16, p<0.0001], and a significant treatment x time interaction [F(23,460)=14.55, p<0.0001] (Figure 2D). Active lever pressing during reinstatement was significantly greater for saline versus metformin treatment in both the first hour (p=0.0171) and the second hour (p=0.0070). Metformin treatment did not significantly modify inactive lever pressing during reinstatement (Figure 2F). These results indicate that metformin in NAcore can reduce cue-induced reinstatement of cocaine seeking in male and female rats (Figure S4).

### Metformin has no effect on maintenance or extinction of cocaine self-administration or spontaneous locomotor activity

The microinjection of metformin into NAcore failed to influence ongoing cocaine self-administration as assessed in a separate subset of female rats. There was no main effect of metformin treatment on inactive lever presses, active lever presses, or infusions during maintenance (Figure S7). Similarly, metformin pretreatment did not impact early extinction lever pressing (Figure S7). As a control experiment, locomotor activity was monitored following saline or metformin microinjection. Intracranial metformin treatment did not alter spontaneous locomotor activity in male rats [main effect of time [F(1.470,11.76)=6.258, p=0.0197] or female rats [main effect of time F(2.029,20.29)=4.676 p=0.021] tested immediately after intracranial infusion, and there were no sex differences observed in the effects of metformin on locomotor activity (Figure S5C, D). Under saline conditions, male and female rats displayed similar levels of locomotor activity and equivalent locomotor habituation (Figure S5A, B). Given that metformin’s effects on cue-induced reinstatement of cocaine seeking were only apparent during the second hour of reinstatement, in separate male subjects we examined the effects of intracranial metformin on locomotion allowing a 1-hour home cage incubation time. In this experiment, metformin was similarly without effect on locomotor activity [main effect of time F(1.645,6.578)=7.86, p=0.0205]; although, locomotion was notable low in the experimental (metformin) and control (saline) group across the entire 45-minute session.

### Sucrose self-administration in male and female Sprague Dawley rats

The experimental timeline is depicted in Figure 4A and mirrors the cocaine experiment. Male and female rats were trained to self-administer sucrose on a FR1 schedule for the first 5 days with the requirement increasing to FR3 for the last 5 days (Figure 4B, 4C). Active and inactive lever pressing for sucrose self-administration differed between sexes with female rats showing higher active lever pressing [F(1,14)=5.786, p=0.0305] and lower inactive lever pressing [F(1,14)=16.64, p=0.0011] compared to male rats (Figure S3C, D). When comparing discrimination index between male and female rats, there was a main effect of session [F(4.352,52.64)=4.105, p=0.0046], a main effect of sex [F(1,14)=20.79, p=0.0004], and a significant sex x session interaction [F(21,254)=2.340, p=0.0011] (Figure 4E). There was a tendency for female rats to maintain higher levels of lever discrimination throughout self-administration and extinction. During self-administration, there was a main effect of time [F(2.510, 34.03)=4.377, p=0.0143] and a main effect of sex [F(1,14)=13.36, p=0.0026] on sucrose pellets administered per day (Figure 4D). Female rats earned more sucrose pellet rewards than male rats (31.20±5.6 versus 57.51±4.9). Akin to cocaine seeking, there was no effect of estrous cycle stage on active lever pressing for sucrose (Figure 4F) or pellets earned (Figure 4G) during the self-administration phase. Extinction day 1 active lever pressing did not differ by estrous cycle but was higher in female rats compared to male rats (p=0.0043) consistent with overall higher active lever pressing throughout sucrose self-administration (Figure 4H).

**Figure 4.**
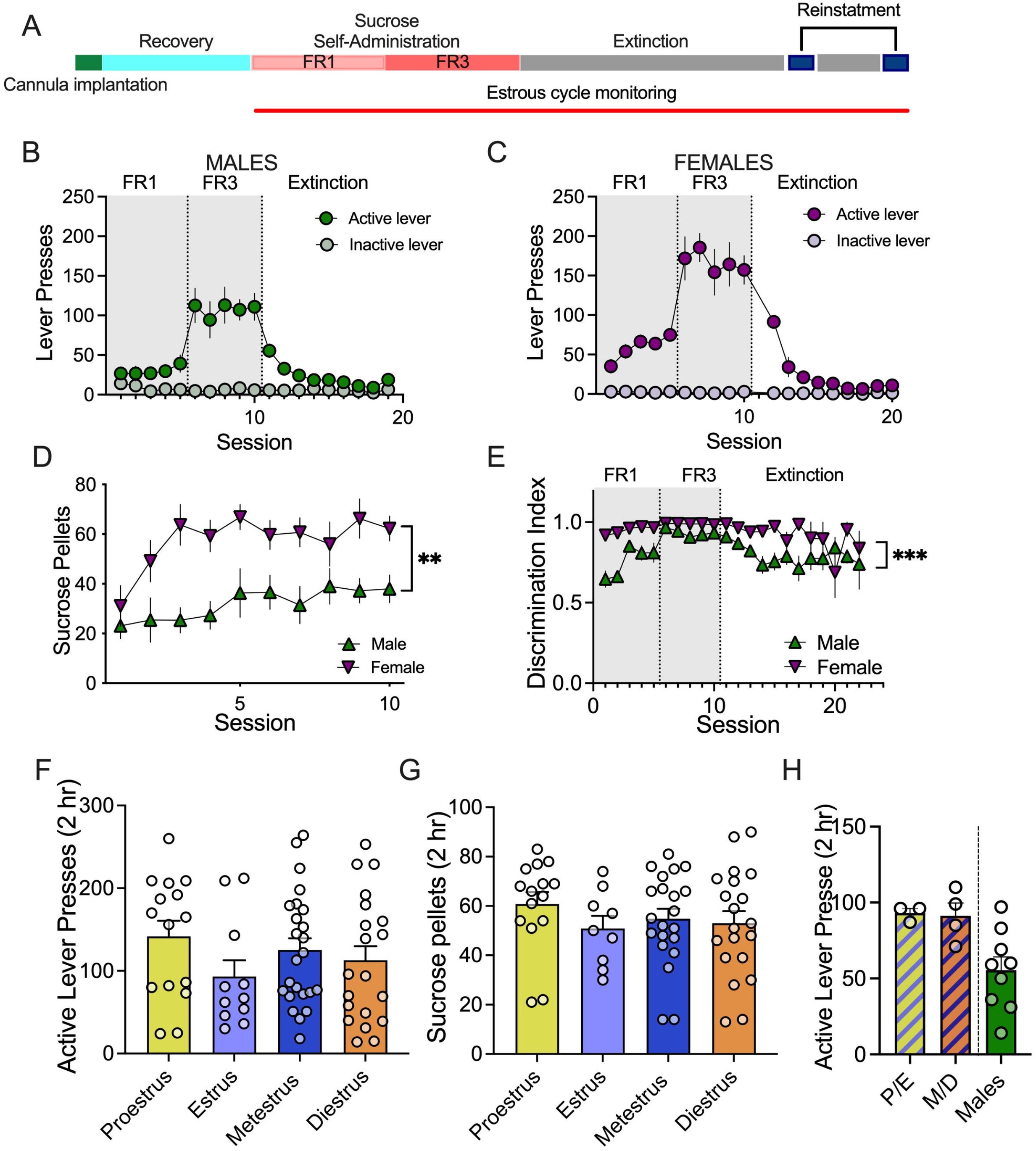
Sucrose self-administration in male and female Sprague Dawley rats. A. Experimental timeline for sucrose self-administration, extinction, and reinstatement. B. Active and inactive lever pressing in males. C. Active and inactive lever pressing in females (mean ± SEM). D. Daily sucrose pellets earned over the 10 days of self-administration was higher in female rats compared to male rats, **p<0.05, n=7-10. E. The discrimination index was higher in female rats compared to males, ***p<0.001, n=7-10 per sex. F. Self-administration active lever pressing did not vary by estrous cycle stage. G. Sucrose pellets delivered during self-administration did not vary by estrous cycle stage. In F. and G. each data point represents the rat mean for each cycle stage. H. Active lever pressing on extinction day 1 did not differ between groups, n=5-7 per phase. P/E = proestrus/estrus and M/D = metestrus/diestrus. All data expressed as mean ± SEM.

### Effects of metformin on cue-induced reinstatement of sucrose seeking

Cue-induced reinstatement of sucrose seeking was performed to query the specificity of the metformin effect for cocaine reward. There was a main effect of lever [F(1,16)=50.54, p<0.0001], a main effect of condition [F(1.573,18.87)=6.941, p=0.0084], and a significant interaction [F(2,24)=7.042, p=0.0039] for cue-induced reinstatement of sucrose seeking in male rats (Figure 5A). Tukey’s multiple comparisons testing indicated a significant cue-induced reinstatement of active lever pressing compared to extinction after both saline (p=0.0093) and metformin microinjection (p=0.0382) with no difference in sucrose seeking between saline and metformin pretreatment conditions (p=0.9998). There was no difference in cumulative active (Figure 5C, [main effect of time F(2.080,27.05)=25.67, p<0.0001] or inactive lever responses (Figure 5E [main effect of time F(2.690,34.97)=15.67, p<0.0001)]) during reinstatement between saline and metformin treatment in male rats. In female rats, there was a main effect of lever [F(1,12)=16.78, p=0.0015], a main effect of condition [F(1.248,14.97)=8.193, p=0.0087], and a significant interaction [F(2,24)=8.113, p=0.002] for cue-induced reinstatement of sucrose seeking (Figure 5B). Tukey’s multiple comparisons testing indicated a significant increase in reinstatement of active lever pressing compared to extinction only after saline microinjection (p=0.0299). When time bin analysis was performed in the female rats, there was a main effect of treatment [F(1,10)=7.989, p=0.018], a main effect of time [F(1.57,15.70)=10.23, p=0.0024], and a significant interaction [F(23,30)=4.016, p<0.0001] (Figure 5D). Metformin treatment did not significantly modify inactive lever pressing during reinstatement (Figure 5F). These results indicate that metformin in NAcore reduces cue-induced reinstatement of sucrose seeking in female but not male rats.

**Figure 5.**
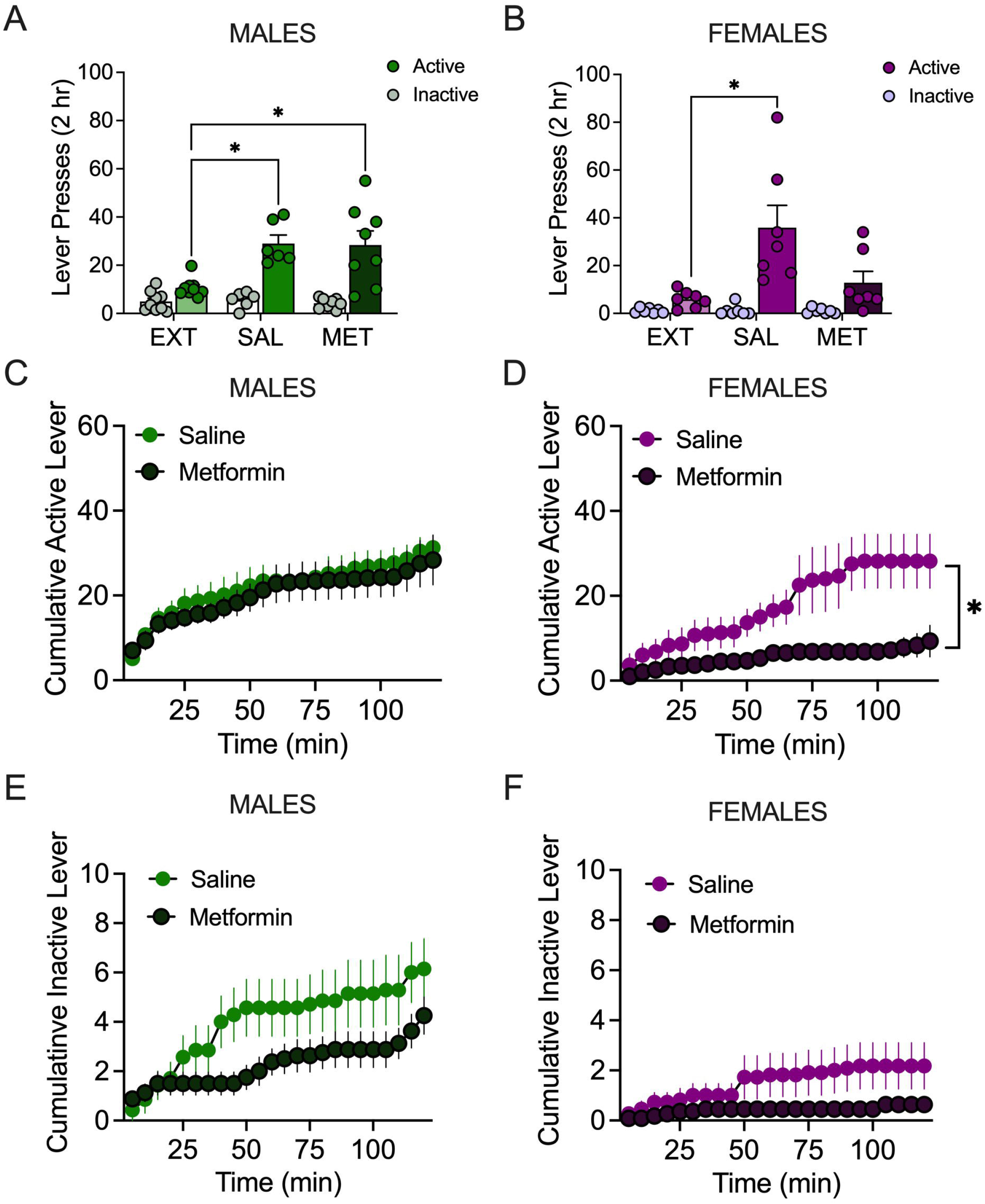
Effects of metformin on cue-induced reinstatement of sucrose seeking. A. Number of active and inactive lever presses during extinction or induced by sucrose-associated cues during reinstatement in male rats, n= 10. *p<0.05 comparing extinction (EXT) to saline (SAL) and metformin (MET). B. Number of active and inactive lever presses during extinction or induced by sucrose-associated cues during reinstatement in female rats, n= 7. *p<0.05 comparing saline to extinction. C. Cumulative active lever pressing in 5-minute bins during cue-induced reinstatement in male rats. D. Cumulative active lever pressing in 5-minute bins during cue-induced reinstatement in female rats. *p<0.05 comparing saline to metformin. E. Cumulative inactive lever pressing in 5-minute bins during cue-induced reinstatement in male rats. F. Cumulative inactive lever pressing in 5-minute bins during cue-induced reinstatement in female rats. All data expressed as mean ± SEM.

## Discussion

Here we show that metformin directly microinjected in NAcore reduces cue-induced reinstatement of cocaine seeking in male and female rats. Effects of intracranial metformin on cocaine seeking were independent of effects on locomotor activity. Diverging from expectations, we found a sex-dependent treatment effect whereby metformin selectively reduced cue-induced cocaine seeking in male rats but generalized to also reduce cue-induced sucrose seeking in female rats. Along with this treatment x sex effect, we report other sex differences in cocaine and sucrose self-administration behavior consistent with sex differences in the reward system.

Metformin and similar biguanides have been used clinically for their blood glucose lowering effects for decades but have recently garnered interest for numerous other purposes including treating cancer, cardiovascular disease, obesity, neurodegeneration, and neurocognitive disorders.^7,21^ Despite long standing use, the mechanism of action of metformin is not fully elucidated. At biologically relevant doses, metformin promotes formation of the AMPK heterotrimeric complex facilitating phosphorylation of the alpha subunit and stimulating kinase activity by >100-fold.^22^ The effects of metformin in the central nervous system have been understudied compared to its peripheral actions. Similarly, the role of AMPK and other cellular energy sensors in important neural processes like learning and memory with clear relevance to reward-related behavior has only recently begun to be appreciated.^23^ Therefore, AMPK activation offers a promising mechanism for metformin effects on reward processing.

One limitation of our current investigation is that we did not confirm the AMPK-dependence of the actions of metformin in our model. Gao and colleagues demonstrated a reduction in pAMPK and an increase in pERK1/2 protein in NAcore following a cocaine self-administration and extinction paradigm analogous to ours.^9^ Infusion of AICAR in the core increased pAMPK and decreased pERK1/2.^9^ We observed a similar pattern with intracranial metformin delivery (Figure S2). Nevertheless, we certainly cannot exclude the possibility that the effects of metformin on cue-induced reinstatement observed here are independent of AMPK activation and instead rely on one of the other purported mechanisms downstream of metformin which we discuss in greater detail below. Future studies will be required to interrogate the role of AMPK and effects of metformin on cocaine-induced neuroadaptations in NAcore that may explain our results.

Our primary finding is that metformin reduces cue-induced cocaine seeking. Relapse-like behavior is governed by dopaminergic and glutamatergic mechanisms,^24^ although there is surprisingly limited direct evidence of a role for dopamine signaling in NAcore in response-contingent cue-induced reinstatement. One study demonstrated that a D1 antagonist and full agonist dose-dependently reduced cue-induced reinstatement for cocaine, while at low doses the agonist also independently promoted reinstated responding in the absence of cues.^25^ Passive presentation of a cocaine-conditioned stimulus (CS) elicits dopamine release in the NAc.^26^ In contrast, response-contingent presentation of a CS after cocaine self-administration, akin to the experimental conditions here, fails to augment NAc dopamine release.^26,27^ Both conditions can facilitate cocaine seeking behavior. These data may imply some level of dissociation between NAc dopamine and various forms of cue-related reinstatement. Systemic metformin increases striatal dopamine and is protective against neurotoxin-induced dopamine depletion^28,29^, but see^30^. Conversely, selective AMPK activation in midbrain slice preparations potentiates K-ATP currents in presumptive dopamine neurons in ventral tegmental area and substantia nigra, an action expected to inhibit neuronal excitability and dopamine release.^31^ *If* dopamine modulation is involved in the effects of our metformin microinjections, the latter mechanism (i.e. decreased dopamine release) could be implicated via metformin actions at dopamine terminals.

The cue-induced reinstatement model of relapse employed here is believed to rely on glutamate transmission. Cocaine self-administration and extinction disrupts glutamate homeostasis in NAcore including decreased basal glutamate levels, decreased glutamate uptake by glial glutamate transporter (GLT-1/EAAT2), and reduced presynaptic mGluR2/3 signaling.^32,33^ The net effect is increased spillover of synaptic glutamate in NAcore during cue-induced reinstatement.^34^ Metformin may be able to correct some of these maladaptations in glutamatergic transmission. For example, repeated metformin reduced elevated glutamate transmission in CA1 pyramidal neurons in a mouse model of LPS-induced depression.^35^ In a genetic mouse model of Fragile X syndrome, chronic metformin treatment improved behavioral deficits and corrected aberrant dendrite morphology and excitatory synaptic activity in hippocampus.^36^ Similarly, metformin may be able to restore glutamate homeostasis in NAcore following cocaine use which would reduce cue-induced reinstatement by limiting glutamate overflow. Alternatively, metformin could reduce cocaine seeking through its actions of reducing oxidative stress^37,38^ although see^39^. Cocaine promotes oxidative stress, including in the ventral striatum, and antioxidant treatments can reduce cue-induced reinstatement of cocaine seeking.^33^ Additional mechanistic studies will be required to disentangle which of these mechanisms (e.g. dopaminergic, glutamatergic, antioxidant, or other) is implicated in the cocaine reinstatement-regulating effects of metformin.

In our experiments, we observed several interesting behavioral sex differences. In humans, females tend to consume cocaine more frequently than males and progress faster from onset of cocaine use to treatment entry.^1,4^ Males have greater difficulty maintaining abstinence following treatment.^40^ Neurobiological mechanisms like sexually dimorphic brain organization or drug-dependent structural adaptations contribute to these differences.^4^ In our cocaine self-administration experiments, the number of infusions earned did not differ by sex, but female rats consumed a higher quantity of cocaine given their higher weight-adjusted unit dose. When self-administering the same unit dose, females correspondingly deliver more cocaine infusions.^41^ We also observed sex differences in cue-induced reinstatement of cocaine seeking with female rats displaying higher levels of active lever pressing compared to males following saline pretreatment (Figure S4A), although the overall pattern of responding across the 2-hour session was similar between sexes indicating similar within-session extinction (Figure S4C). Consistent between cocaine and sucrose self-administration experiments, female rats maintained higher lever discrimination during extinction which could represent a difference in motivation that emerges in the absence of reinforcement independent of initial reinforcer.

Preclinical literature suggests that females may be more sensitive to natural rewards than males. Female rats showed a higher preference for chambers paired with palatable food (high fat, high sugar)^42^ and higher demand for self-administering palatable food rewards at null cost compared to male rats.^43^ Agreeing with this literature, our female rats self-administered more sucrose rewards and displayed higher active lever pressing throughout sucrose self-administration (Figure S3C, D). Although our sucrose self-administration and cue reinstatement was designed as a control study to determine whether metformin administration selectively decreases cue-induced reinstatement of cocaine seeking, ultimately this experiment garnered some of the most interesting results. We found a sex-dependent treatment effect whereby metformin reduced cue-induced sucrose seeking in female but not male rats. This could suggest an increased sensitivity to metformin’s effects on reward seeking in females. It will be important to determine if metformin dosage can be calibrated in females to mitigate the risk of producing anhedonia. Alternatively, these results may speak more to sex differences in sucrose metabolism and/or reward processing between male and female rats related to the differential behavioral sensitivity for palatable food described above.^42,43^ It is not completely clear if or how AMPK activation might be involved. One study showed that female rats were more sensitive than males to the detrimental metabolic effects of chronic fructose, and AMPK was selectively activated in the liver of fructose-fed female rats.^18,44^ While our cocaine reinstatement results suggest cocaine similarly regulates AMPK activity in the NAcore in males and females^9^, sucrose self-administration may produce divergent neurochemical changes that have yet to be investigated. ICV metformin inhibits food intake, but this effect is thought to be mediated through inhibition of neuropeptide Y in the hypothalamus.^18^ The hypothalamus regulates homeostatic feeding while the nucleus accumbens modulates palatable food intake.^45,46^ The role of extra-hypothalamic AMPK in both homeostatic and hedonic feeding mechanisms thus warrants further exploration.

The pharmacodynamics of metformin do not differ by sex in healthy control subjects,^11^ but there are multiple examples of metformin producing sex-specific effects under certain conditions.^47,48^ For example, metformin has more beneficial effects on glucose metabolism in diabetic males compared with females.^49^ Mechanistic links exist between metformin, AMPK, and sex steroid hormones. Sex differences were detected in expression of the organic cation transporter (OCT2), one of the transporters responsible for moving metformin across the cellular membrane.^50^ Higher mRNA expression was detected in cortex and hippocampus of female compared to male mice although no sex differences were observed at the protein level.^50^ Estrogen lowers the activity of OCT2 in males while testosterone increases OCT2 activity in both males and females in the rat kidney.^51^ It is unknown if similar hormonal regulation of OCT2 (or other OCT family members) is present in the brain. If present, this could significantly impact metformin’s effects on neurobiological processes. OCT2 immunoreactivity is observed at high levels in the rat NAc and demonstrates significant overlap with staining for the vesicular acetylcholine transporter (VAChT) and dopamine transporter (DAT).^52^ Functionally, OCT2 participates in the clearance of serotonin and norepinephrine in the mouse striatum.^53^ Taken together, the potential dual role of this transporter in regulation of neurotransmission and mediation of the pharmacologic response to metformin make this protein an interesting target for future investigation.

We might have predicted an interaction between estrous cycle stage and metformin treatment for cue-induced reinstatement given that 17⊓-estradiol (E2) influences the activation of AMPK.^54^ An analysis of estrous cycle dependent effects associated with relapse to cocaine seeking demonstrates a main effect of treatment (Figure S4D, [F(1,18)=9.922, p=0.0055]) but no main effect of estrous cycle stage or interaction. Nevertheless, Sidak’s multiple comparisons indicate that metformin’s ability to reduce cue-induced reinstatement of cocaine seeking in females was driven by effects observed during diestrus/metestrus. Few other estrous cycle stage dependent effects emerged within the data. We observed significantly higher active lever pressing during the first cocaine extinction session when female rats were in proestrus/estrus opposed to metestrus/diestrus (Figure 1H). The follicular phase (proestrus/estrus) is characterized by an increase in estrogen levels and a decrease in progesterone levels. This is notable because estrogen has been found to play a role in increasing the reinforcing effects of psychostimulants.55 In contrast, progesterone which peaks in the luteal phase attenuates positive reinforcing effects of psychostimulants.56 Interestingly, we did not find any estrous cycle-dependent difference in active lever pressing during the first extinction session after sucrose experience. We also did not have enough observations to detect significant differences during cue-induced reinstatement of sucrose seeking.

Cocaine use disorder remains without any available pharmacotherapy. These data represent a first step in the process of investigating metformin for its potential in treating cocaine use disorder. We were encouraged by the observation that intracranial metformin reduces cue-induced reinstatement after cocaine self-administration and extinction. In future studies, it will be important to extend these data to other reinstatement modalities (context, cocaine-primed, stress-induced) because while these models share important circuit mechanisms they do not completely overlap.57 These differences may inform the precise circumstances under which metformin may be most effective as a therapeutic option. Similarly, here we only tested intracranial administration of metformin focusing on a site of action in the NAcore due to its well-established role in cue-induced reinstatement. This strategy is impractical for human translation; therefore, it will be necessary to determine if systemic metformin delivered acutely or chronically can likewise alter cocaine seeking. Intracranial metformin produced sex-dependent effects on cue-induced sucrose seeking, reducing reward seeking in females but not males. Additional studies are warranted to parse the effect on natural reward. Given the pleiotropic effects of metformin in the whole body and brain, we will endeavor to ultimately elucidate which speculative molecular mechanism accounts for the therapeutic benefits in our model. Moreover, there may be potential for extending this pharmacotherapy to other amphetamine-like psychostimulants given the prominent effects of these drugs on brain energy metabolism.

## Acknowledgements

This work was supported by startup funds from the University of Minnesota and the Medical Discovery Team on Addiction and the National Institutes of Health (R00 DA0411462 to SS and T32 DA007234 to MS).

## Authors contribution

AC and LW performed the experiments, analyzed the data, and wrote the manuscript. SM, NI, AS, and MS performed experiments. SS was responsible for the study concept and design, supervised the experiments, analyzed the data, wrote the manuscript. All authors approved the final version for publication.

## Data Availability Statement

The data that support the findings of this study are available from the corresponding author upon reasonable request.

