## supplementary materials for "Metformin in nucleus accumbens core reduces cue-induced cocaine seeking in male and female rats"

**Supplementary Information**

**Expanded Methods**

**Subjects**: 137 male and female Sprague Dawley rats (225-275g on arrival) were purchased from Envigo Laboratories for these studies. 109 subjects were included in the final dataset. 16 rats were excluded for failing to acquire or extinguish behavior (12%); 12 rats were excluded due to occluded cannula or missed placements (9%). Rats were acclimated to the facilities for 5-7 days prior to surgery. Subjects were double housed in individually ventilated cages in a temperature (70-71℉) and humidity (38-46%) controlled holding facilities. Standard chow and water were available *ad libitum* with the exception of the 24-hour period preceding a single food training session performed prior to cocaine self-administration. Animals were kept on a 14 /10 hour light-dark cycle with all experiments conducted during the light phase. Procedures were preapproved by the Institutional Animal Care and Use Committee (IACUC) of the University of Minnesota and follow National Institute of Health (NIH) guidelines.

**Intracranial and Intravenous Surgery**: Rats (250-300g at time of surgery) were anesthetized with vaporized isoflurane (2-5%). Carprofen (5 mg/kg, i.p.) was provided for peri- and post-operative analgesia, and ceftriaxone (200 mg/kg, i.m.) was administered as a prophylactic and post-surgical antibiotic. Rats were implanted with intravenous jugular catheters for cocaine self-administration. The silastic catheter tubing, secured to the right jugular vein with silk sutures, was passed subcutaneously to the middle of the back where it terminated in a connector consisting of a modified 22-gauge cannula (Plastics One, Roanoke, VA) embedded in dental cement attached to surgical mesh (Atrium, Hudson, NH). Stereotaxic surgery (Kopf Instruments, Tujunga, CA) was performed to deliver bilateral guide cannula (Plastics One) to the NAcore, coordinates ±1.8 mm ML, +1.5 mm AP, -5.5 mm DV at a 0° angle.^1^ Coordinates were slightly modified for the experiments illustrated in Supplementary Figure S7: ±1.6 mm ML, +2.3 mm AP, -5.5 mm DV at a 0° angle. Cannulae were affixed to the skull using acrylic dental cement stabilized with 2-3 jeweler’s screws. Metal obturators were placed in the cannula to block debris. Post-operative care was provided for 72 hours after surgery. Catheter patency was confirmed prior to initiating cocaine self-administration with intravenous administration of 0.05-0.1 mL xylazine (5mg/mL). Loss of muscle tone within 5 seconds indicated satisfactory catheter patency. Rats without patent catheters were re-catheterized in the opposite (left) jugular vein.

**Drugs:** Cocaine hydrochloride was purchased from Boynton Pharmacy at the University of Minnesota. Cocaine solution (4 mg/ml) was prepared in sterile 0.9% saline and delivered at a volume of 0.05 ml/infusion. The average mg/kg/infusion for male and female rats was 0.59 and 0.74 mg/kg, respectively over the course of self-administration (Figure S1). Metformin hydrochloride (ApexBio, Sigma-Aldrich), 1,1-dimethylbiguanide) was prepared at 250 μg/μl in sterile water, and 125 μg/hemisphere was microinjected into NAcore. The metformin dose was chosen based on published studies of intracerebroventricular (ICV) administration in rats using doses ranging from 3 μg to 1 mg.^2–5^ Metformin (200 μg, ICV) reduced food intake in male Sprague Dawley rats, an effect associated with increased phospho-AMPK in hypothalamus indicative of enzyme activation.^2^ Metformin’s effects on food intake were observed up to 24 hours after microinjection. After an acute oral administration of 150 mg/kg metformin, brain levels of metformin measured by high performance liquid chromatography peaked at 6 hours post-administration.^6^ Metformin levels did not fall below the limit of detection within the 48-hour measurement period.^6^ Thus, metformin is expected to be bioavailable for the duration of the two-hour tests. Saline or metformin was microinjected in NAcore with brain dissections collected after 1 hour in a subset of animals. As expected, metformin increased levels of phospho-AMPK and showed a trend toward decreased phospho-ERK consistent with AMPK activation (Figure S2).

**Cocaine and sucrose self-administration, extinction, and cue-induced reinstatement**: Rats underwent one 2-hour session of operant training for food pellets in the absence of cues prior to beginning daily 2-hour cocaine self-administration sessions. Rats experienced a single overnight food restriction preceding the food training session. Training was completed in standard operant boxes (MedAssociates) equipped with a house light, multiple tone generator, two retractable levers, two stimulus lights, food trough, and pellet dispenser. Rats self-administered cocaine on a fixed ratio 1 (FR1) reinforcement schedule. Each cocaine infusion was paired with a discrete white light and tone dual stimulus (5 second, 2900 Hz, 80 dB) followed by a 15 second timeout to prevent cocaine overdose. Inactive lever presses were recorded but had no consequences. Catheters were flushed daily with heparinized saline (100 USP units/mL i.v.) to maintain patency throughout self-administration. Rats were required to reach a criterion of 10 days of self-administration with at least 10 cocaine infusions. Next, subjects transitioned to extinction (minimum 7 days) whereupon pressing the formerly active lever no longer resulted in delivery of cocaine or cues. Extinction occurred until active lever pressing was reduced to less than or equal to 30% of the mean established during self-administration. Cue-induced reinstatement followed extinction training, whereupon pressing of formerly active lever resulted in presentation of drug-paired cues but no drug delivery during a 2-hour session. The protocol for sucrose self-administration was analogous to that used for cocaine, described above. Rather than cocaine infusions, sucrose pellets (45 mg, BioServ) were paired with a discrete light and tone cue. Rats were trained to self-administer sucrose pellets for at least 5 days on an FR1 reinforcement schedule and then progressed to 5 days on an FR3 reinforcement schedule with acquisition criterion of ≥10 sucrose pellets earned per session. Extinction and cue-induced reinstatement conditions were identical to cocaine self-administration.

**Locomotor Activity:** Spontaneous locomotor activity was recorded with a USB webcam placed over the test chamber (60cm x 60cm square acrylic box) for 45 minutes. Videos were analyzed for total distance traveled using ezTrack video analysis software.^7^ Animals were given microinjections of saline or metformin to the NAcore immediately before or 1 hour prior to placement in test chamber.

**Vaginal Cytology:** Vaginal cytology samples were collected from female rats daily.^8^ Saline (150 μL) was gently flushed 2-3x into the vaginal opening. The sample was spread onto a glass slide, left to air-dry, stained with a differential stain kit (NewcomerSupply), and imaged at 10x. Samples were compared to published examples of vaginal cytology and categorized into stages using previously published criteria.^8^ The proestrus stage is identified by a predominance of nucleated epithelial cells. Estrus was defined as predominantly cornified epithelial cells. Metestrus cytology has roughly equal numbers of cornified epithelial cells and leukocytes. Composition of cells during diestrus is >75% leukocytes, <25% epithelial cells.

**Western blotting:** Animals were rapidly decapitated and tissue samples micro-dissected for sample preparation. Whole-cell lysates were prepared in RIPA buffer supplemented with 1% SDS and 1x protease and phosphatase inhibitors. Protein concentration was determined using the Rapid Gold BCA Protein Assay (Pierce). Electrophoresis was performed according to standard protocols using 4-12% Criterion XT Bis-Tris precast gels (Bio-Rad) run in XT-MOPS buffer (Bio-Rad) at 180 V for 1 hour.^9^ Gels were transferred to nitrocellulose membranes using the Trans-Blot Turbo transfer system (Bio-Rad). Primary antibodies were used as follows: phospho-AMPKɑ Thr172 (Cell Signaling 2535, 1:500), AMPKɑ (Cell Signaling 2532, 1:1000), phospho-p44/42 MAPK (ERK1/2) Thr202/Tyr204 (Cell Signaling 4370, 1:2000), p44/42 MAPK (ERK1/2) (Cell Signaling 9102, 1:1000), GAPDH (Cell Signaling 2118, 1:5000). HRP-conjugated secondary antibodies (Cell Signaling 7074 and 7076) and Super Signal West Dura ECL reagent (ThermoFisher) were used for chemiluminescent detection. Blots were imaged on the iBright FL1500 (ThermoFisher).

**Statistical Analysis:** GraphPad Prism version 9.0.1 (San Diego, California USA) was used to perform statistical analysis. For comparing multiple measurements in the same experiment, the data were analyzed using 1- or 2-way ANOVAs or mixed effects models as appropriate for each experiment. Mixed effect models were utilized when data were missing at random (e.g., due to COVID-19 research restrictions) and a full repeated measures ANOVA could not be run. Tukey’s or Sidak’s *post-hoc* testing was applied for multiple comparisons and *p*<0.05 was considered statistically significant. Data are presented as mean ± standard error of the mean (SEM). To evaluate metformin’s effects on reinstatement in male or female rats, summary data were analyzed by two-way ANOVA using within-subjects comparisons using lever (active vs inactive) and condition/test (extinction, saline reinstatement, or metformin reinstatement) as factors. For time course experiments, data were analyzed by two-way ANOVA using time and treatment as factors. To examine sex differences, data were analyzed by two-way ANOVA using sex as a between-subject factor and time (time, day, or phase of experiment) as a repeated factor. Estrous cycle stage comparisons were performed using one-way ANOVAs. When only two groups were compared, a Student’s t-test was used. The discrimination index between active and inactive lever pressing during self-administration and extinction was determined using the equation: (Active)/(Active + Inactive).

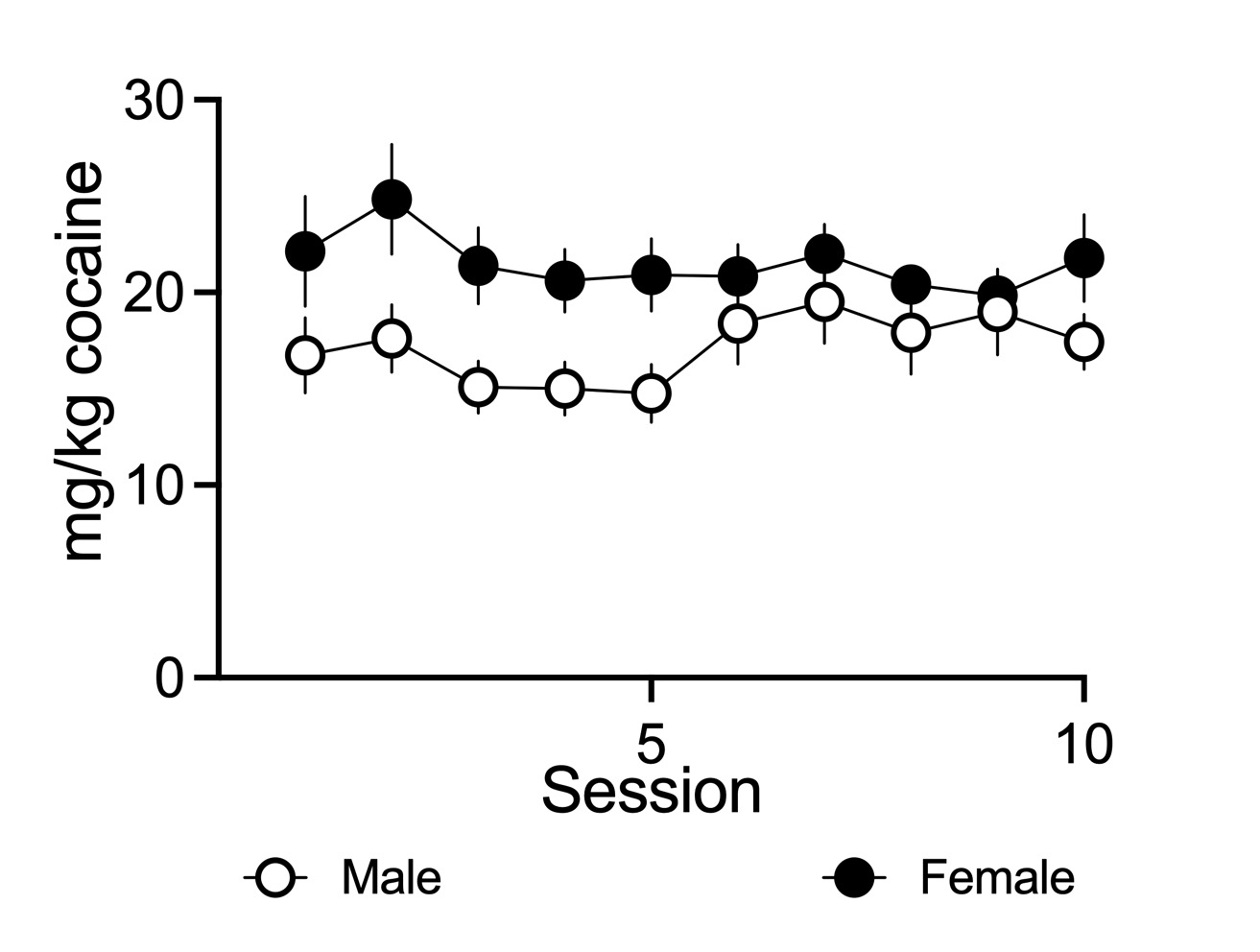

**Supplementary Figure 1.** Daily cocaine intake in mg/kg expressed as mean ± SEM over the 10 days of self-administration at criteria compared between male (n=10) and female (n=12) rats. Two-way ANOVA indicates a main effect of sex [F(1,20)=6.735, p=0.0173] for cocaine intake. Females consumed more cocaine on average than male rats across 10 days of self-administration T(20)=2.260, p=0.0351. All data expressed as mean ± SEM.

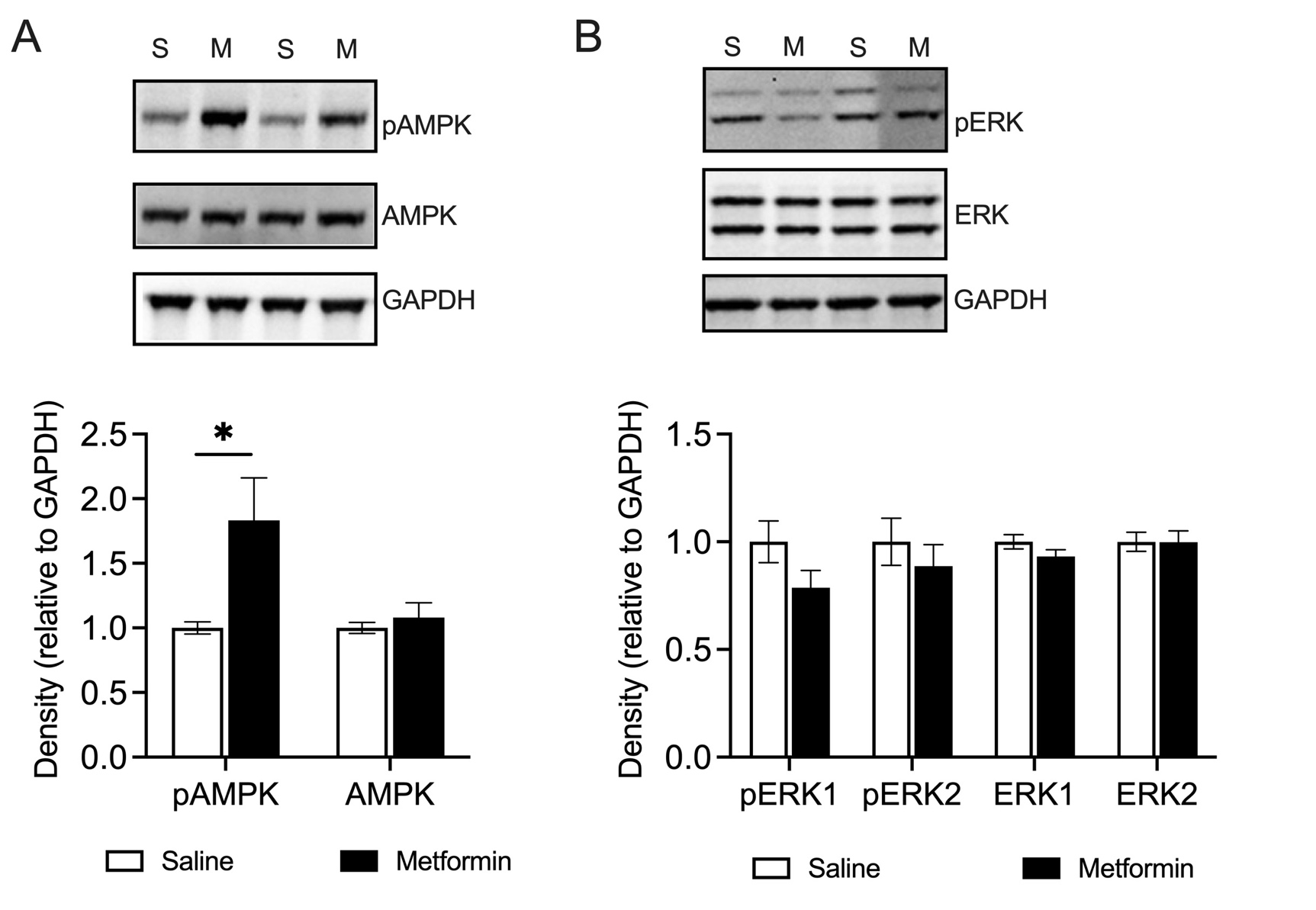

**Supplementary Figure 2.** Western blot analysis of nucleus accumbens core following microinjection of metformin or saline. Tissue dissections were taken one hour after microinjection. A. Metformin increases levels of phosphorylated AMPK but not total AMPK. T(8)=2.502, p=0.0368. B. Metformin did not significantly alter pERK or total ERK levels, but there was a trend toward the expected decrease, p=0.1038.

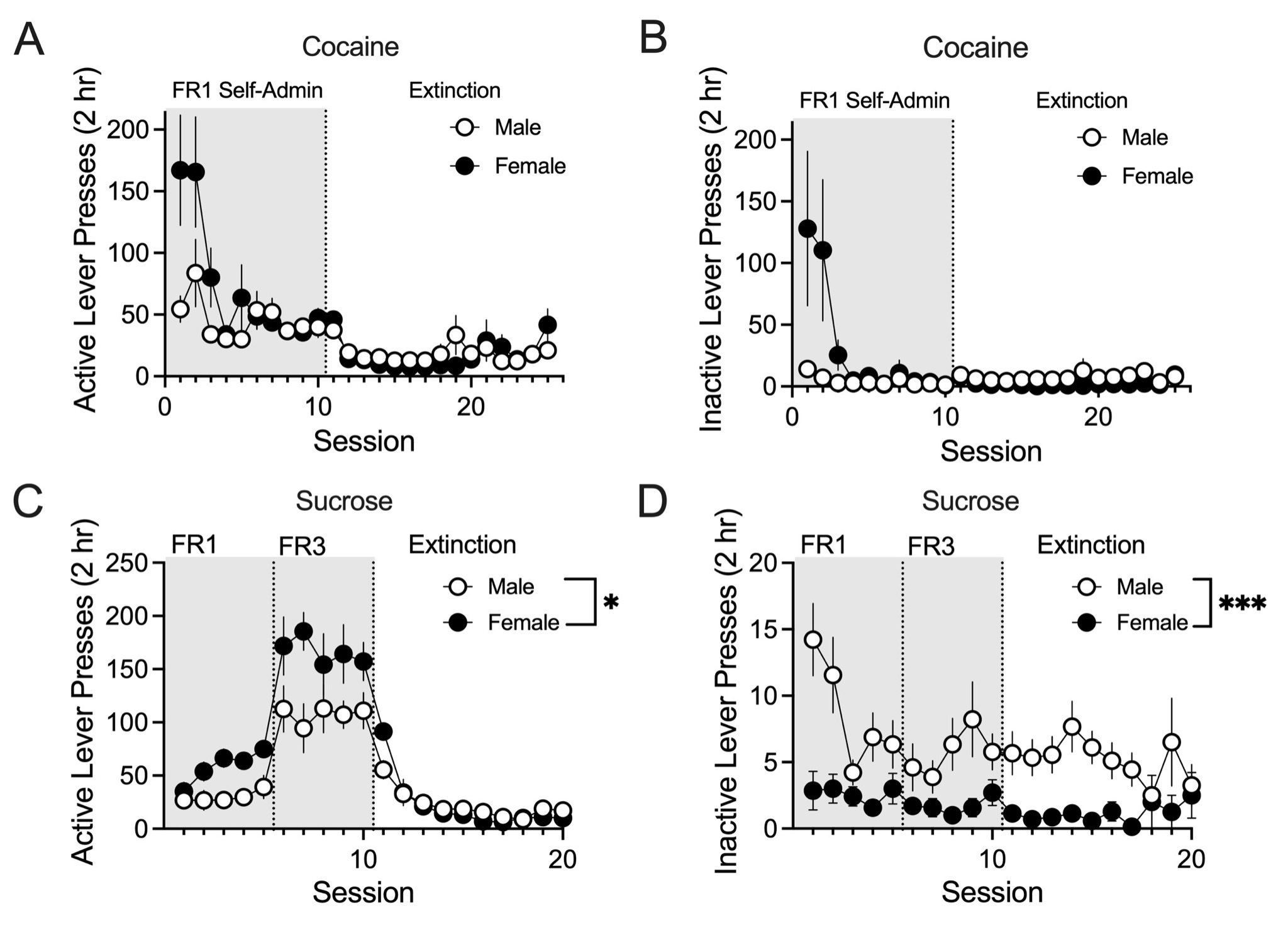

**Supplementary Figure 3.** Comparisons of cocaine and sucrose self-administration behavior analyzed by sex corresponding to data in Figure 1 and Figure 4, n=10-12 per group. A. Active lever pressing during cocaine self-administration and extinction in male and female rats. Mixed-effects analysis indicates a main effect of time [F(2.057,33.34)=9.590, p=0.0005] and a Time x Sex interaction [F(24,389)=2.538, p=0.0001]. B. Inactive lever pressing during cocaine self-administration and extinction in male and female rats. Mixed-effects analysis indicates no significant main effects. C. Active lever pressing during sucrose self-administration and extinction in male and female rats. *p<0.05 comparing males and females. Mixed-effects analysis indicates a main effect of time [F(3.784,46.48)=37.56, p<0.0001], a main effect of sex [F(1,14)=5.786, p=0.0305] and a Time x Sex interaction [F(21,258)=3.344, p<0.0001]. D. Inactive lever pressing during sucrose self-administration and extinction in male and female rats. ***p<0.001 comparing males and females. Mixed-effects analysis indicates a main effect of time [F(4.552,55.93)=2.542, p=0.0429], a main effect of sex [F(1,14)=16.64, p=0.0011] and a Time x Sex interaction [F(21,258)=1.451, p=0.0950]. All data expressed as mean ± SEM.

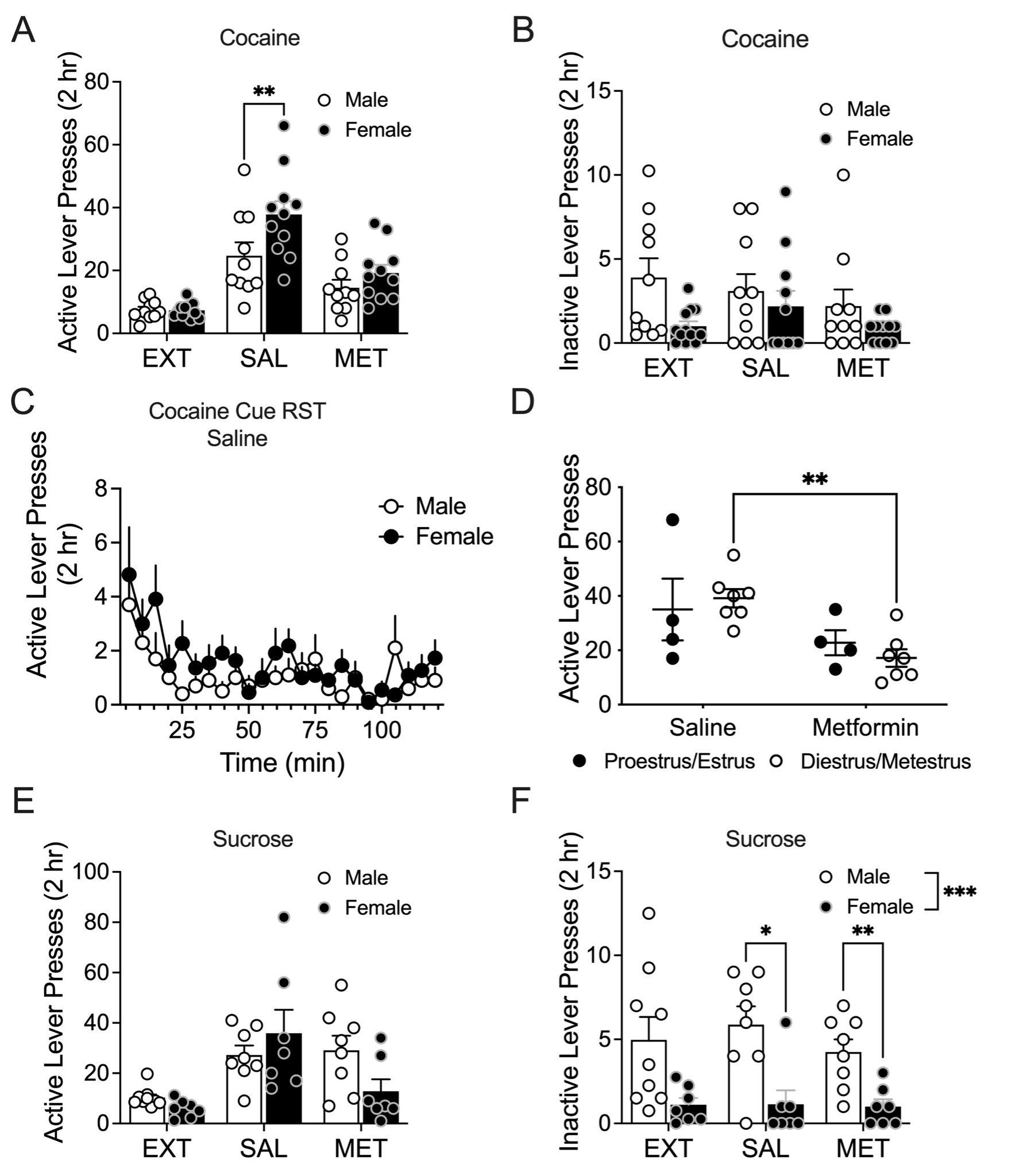

**Supplementary Figure 4.** Comparisons of cocaine and sucrose cue-induced reinstatement behavior analyzed by sex corresponding to data in Figure 2 and Figure 5, n=10-12 per group. EXT = extinction, SAL = saline, MET = metformin. A. Active lever pressing during cue-induced reinstatement of cocaine seeking in male and female rats. Mixed-effects analysis indicates a main effect of condition [F(2,38)=37.85, p<0.0001], a main effect of sex [F(1,204)=5.592, p=0.0283], and no interaction. Sidak’s multiple comparison testing indicates a significant sex difference in cue-induced reinstatement under saline pretreatment conditions (**p=0.0065) with a higher response in females. B. Inactive lever pressing during cue-induced reinstatement (RST) of cocaine seeking in male and female rats. Mixed-effects analysis indicates no significant main effects. C. Time course of active lever pressing during cue-induced reinstatement of cocaine seeking following saline pretreatment in male and female rats. Two-way ANOVA indicates a main effect of time [F(7.283,138.4)=3.604, p=0.0012]. D. Active lever pressing during cue-induced reinstatement of cocaine seeking in female rats separated by estrous cycle stage condition. Two-way ANOVA indicates a main effect of treatment [F(1,18)=9.922, p=0.0055] and Sidak’s multiple comparisons reveal a significant treatment effect selectively for diestrus/metestrus stages (**p=0.0070). E. Active lever pressing during cue-induced reinstatement of sucrose seeking in male and female rats. Mixed-effects analysis indicates a main effect of condition [F(1.499,19.49)=12.27, ***p=0.0008] and Sex x Condition interaction [F(2,26)=3.374, p=0.0498] F. Inactive lever pressing during cue-induced reinstatement of sucrose seeking in male and female rats. Mixed-effects analysis indicates a main effect of Sex [F(1,14)=17.84, p=0.0009]. Sidak’s multiple comparison testing indicates a significant sex difference in inactive lever pressing during cue-induced reinstatement under saline (*p=0.0135) and metformin (**p=0.0096) pretreatment conditions. All data expressed as mean ± SEM.

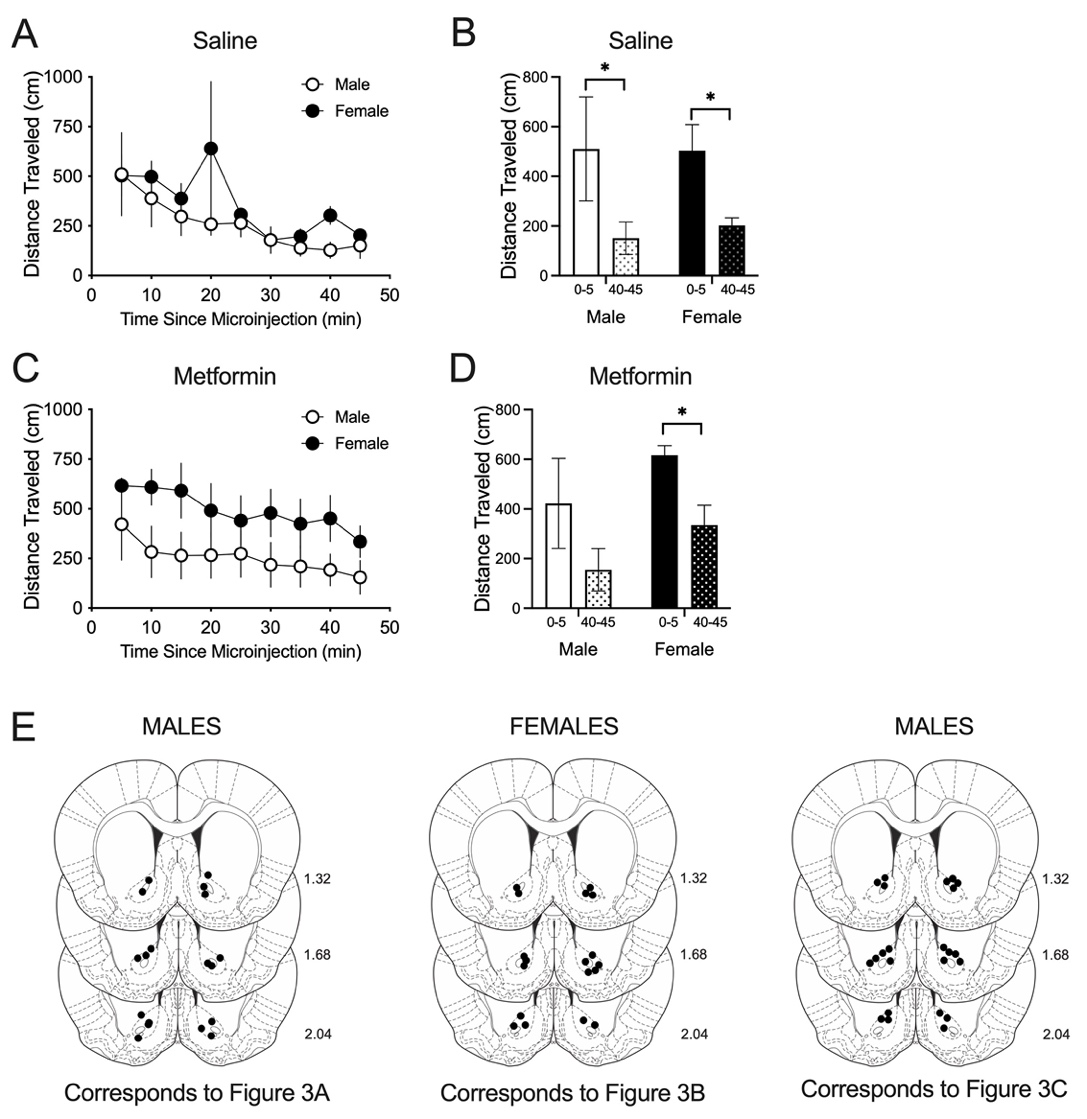

**Supplementary Figure 5.** Comparisons of locomotor activity analyzed by sex corresponding to data in Figure 3, n=5-7 per group. A. Locomotor activity in male and female rats immediately after microinjection of saline. Two-way ANOVA indicates a main effect of time [F(2.052,16.42)=4.040, p=0.0366] but no main effect of sex. B. Comparison of the first and last 5-minute bins of locomotor activity in male and female rats after saline microinjection. Paired t-tests reveal habituation of locomotor activity in male [T(4)=2.913, *p=0.0435] and female [T(4)=3.474, *p=0.0255] rats as evidenced by lower activity during the 40-45 minute epoch as compared to the 0-5 minute epoch. C. Locomotor activity in male and female rats immediately after microinjection of metformin. Two-way ANOVA indicates a main effect of time [F(2.117,21.17)=6.258, p=0.0066] but no main effect of sex. D. Comparison of the first and last 5-minute bins of locomotor activity in male and female rats after metformin microinjection. Paired t-tests reveal habituation of locomotor activity in female rats [T(6)=3.145, *p=0.0199] as evidenced by lower activity during the 40-45 minute epoch as compared to the 0-5 minute epoch. This comparison was not statistically significant in males after metformin pretreatment (p=0.0913). E. Histological verification of cannula placement for animals included in locomotor activity analysis. All data expressed as mean ± SEM.

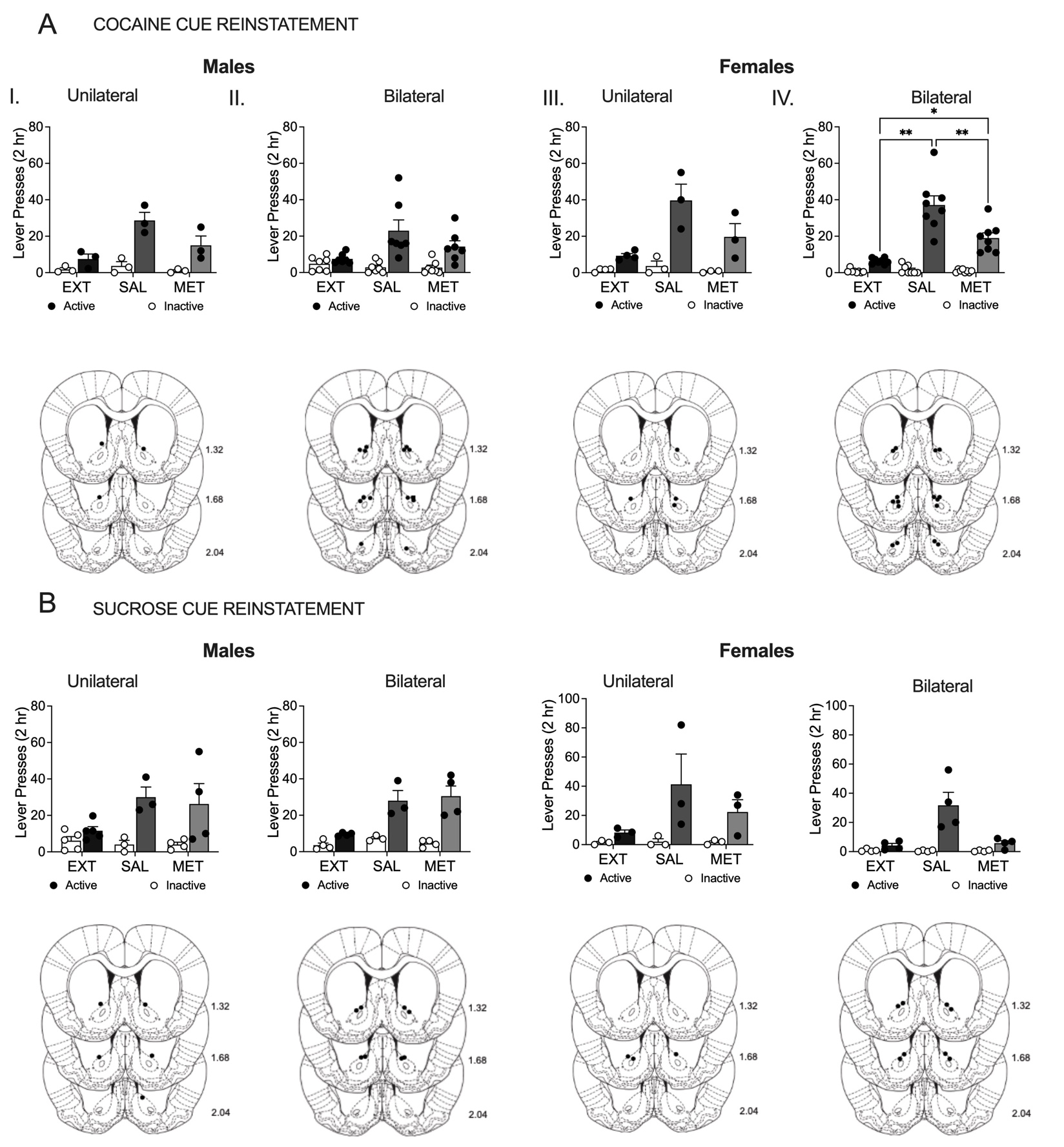

**Supplementary Figure 6**. Effects of metformin on cue-induced reinstatement of cocaine and sucrose seeking with data subdivided to examine unilateral versus bilateral microinjections corresponding to data in Figure 2 and Figure 5. EXT = extinction, SAL = saline, MET = metformin. Histological verification of cannula placement for cocaine and sucrose self-administration experiments indicated below the corresponding summary data. A. Cocaine Cue Reinstatement. I. Number of active and inactive lever presses during extinction or induced by cocaine-associated cues during reinstatement in male rats with unilateral microinjections. Two-way ANOVA indicates a main effect of Lever [F(1,4)=77.23, p=0.0009] but the effect of Condition was not quite significant [F(1.347,5.387)=5.475, p=0.0574]. II. Number of active and inactive lever presses during extinction or induced by cocaine-associated cues during reinstatement in male rats with bilateral microinjections. Two-way ANOVA indicates a main effect of Lever [F(1,12)=23.76, p=0.0004] and a Lever x Condition interaction [F(2,24)=4.405, p=0.0235]. II. Number of active and inactive lever presses during extinction or induced by cocaine-associated cues during reinstatement in female rats with unilateral microinjections. Two-way ANOVA indicates a main effect of Lever [F(1,6)=24.00, p=0.0027], a main effect of Condition [F(1.886,7.544)=10.35, p=0.0073] and a Lever x Condition interaction [F(2,8)=7.729, p=0.0135]. IV. Number of active and inactive lever presses during extinction or induced by cocaine-associated cues during reinstatement in female rats with bilateral microinjections. Two-way ANOVA indicates a main effect of Lever [F(1,14)=68.70, p<0.0001], a main effect of Condition [F(1.572,22.01)=28.16, p<0.0001] and a Lever x Condition interaction [F(2,28)=25.14, p<0.0001]. Tukey’s multiple comparisons testing revealed significant differences in active lever pressing by condition: **p<0.01 comparing saline to extinction or metformin and *p<0.05 comparing metformin to extinction. B. Sucrose Cue Reinstatement. I. Number of active and inactive lever presses during extinction or induced by sucrose-associated cues during reinstatement in male rats with unilateral microinjections. Mixed-effects analysis indicates a main effect of Lever only [F(1,18)=17.06, p=0.0006]. II. Number of active and inactive lever presses during extinction or induced by sucrose-associated cues during reinstatement in male rats with bilateral microinjections. Mixed-effects analysis indicates a main effect of Lever [F(1,6)=3038, p=0.0015], a main effect of Condition [F(1.855,9.273)=10.90, p=0.0041], and a Lever x Condition interaction [F(2,10)=7.464, p=0.0104]. III. Number of active and inactive lever presses during extinction or induced by sucrose-associated cues during reinstatement in female rats with unilateral microinjections. Two-way ANOVA indicates no significant main effects. IV. Number of active and inactive lever presses during extinction or induced by sucrose-associated cues during reinstatement in female rats with bilateral microinjections. Two-way ANOVA indicates a main effect of Lever [F(1,6)=14.60, p=0.0087], a main effect of Condition [F(1.045,6.268)=9.785, p=0.0187] and a Lever x Condition interaction [F(2,12)=10.09, p=0.0027]. All data expressed as mean ± SEM. Overall, we qualitatively observed equivalent trends in these behavioral data with bilateral versus unilateral microinjections for cue-induced reinstatement of cocaine and sucrose seeking in both males and females. This supports combining these conditions for the final in-text analysis.

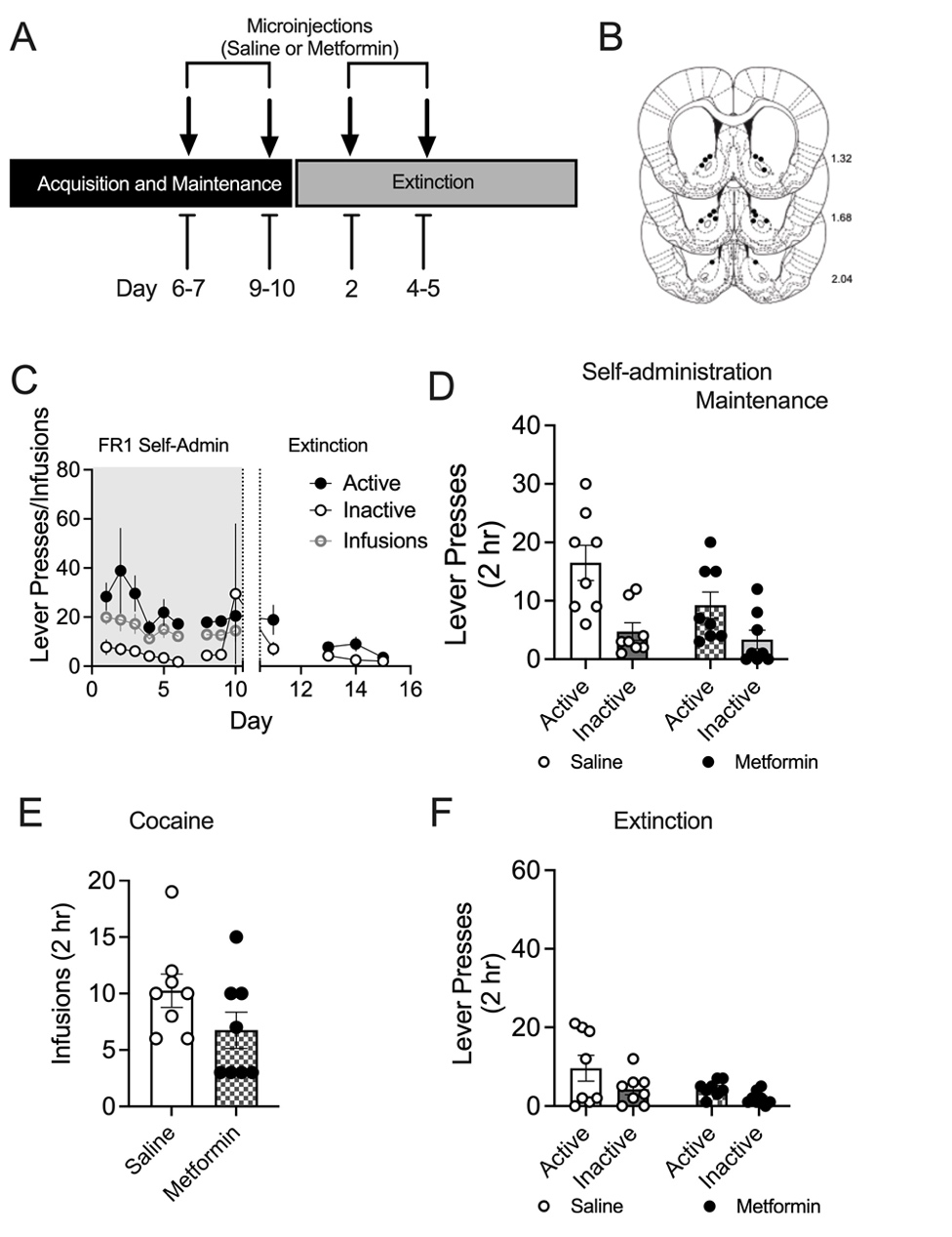

**Supplementary Figure 7.** Effects of metformin on cocaine self-administration maintenance and extinction. A. Experimental timeline for cocaine self-administration and extinction with saline and metformin microinjections. B. Histological verification of cannula placement. C. Active and inactive lever pressing during self-administration and extinction on non-test days. D. Active and inactive lever pressing during cocaine self-administration comparing saline and metformin pretreatment. Paired t-tests for active (T(7)=1.585, p=0.1570) and inactive (T(7)=0.763, p=0.4704) lever pressing reveal lack of metformin effect on maintenance of self-administration. E. Cocaine infusions during cocaine self-administration did not differ between saline and metformin pretreatment (T(7)=1.478, p=0.1829). F. Active and inactive lever pressing during extinction, comparing saline and metformin pretreatment. Paired t-tests for active (T(7)=1.771, p=0.1198) and inactive (T(7)=2.049, p=0.0796) lever pressing reveal lack of metformin effect on extinction of cocaine self-administration. All data expressed as mean ± SEM.
